# Defining the extent of gene function using ROC curvature

**DOI:** 10.1101/2021.09.03.458825

**Authors:** Stephan Fischer, Jesse Gillis

## Abstract

Machine learning in genomics plays a key role in leveraging high-throughput data, but assessing the generalizability of performance has been a persistent challenge. Here, we propose to evaluate the generalizability of gene characterizations through the shape of performance curves. We identify Functional Equivalence Classes (FECs), uniform subsets of annotated and unannotated genes that jointly drive performance, by assessing the presence of straight lines in ROC curves. FECs are widespread across modalities and methods, and can be used to evaluate the extent and context-specificity of functional annotations in a data-driven manner. For example, FECs suggest that B cell markers can be decomposed into shared primary markers (10 to 50 genes), and tissue-specific secondary markers (100 to 500□genes). In addition, FECs are compatible with a wide range of functional encodings, with marker sets spanning at most 5% of the genome and data-driven extensions of Gene Ontology sets spanning up to 40% of the genome. Simple to assess visually and statistically, the identification of FECs in performance curves paves the way for novel functional characterization and increased robustness in analysis.

## Introduction

Characterizing the functional properties of genes across conditions, species, and other perturbations is a central challenge in post genome biology. As data sets increase in size and complexity, exploiting methods from machine learning and AI research has become increasingly valuable to parse vast data collections for subtle convergent signals^1–5^. However, the complexity and customized nature of these methods create interpretation problems of their own. Establishing a consensus framework for comparative evaluation has been essential to progress, often using systematic data resources, and with well-defined performance metrics. In particular, many problems in genomics map to a supervised learning framework with a goal of determining functional sets of genes from partial annotations and feature data. A correspondingly high number of methods and assessments report comparative evaluation using traditional machine learning statistics, such as the area under the receiver-operator characteristic curve (AUROC). However, genomics poses unique challenges and opportunities relating to its unusual scalability, both across novel contexts (conditions or species) and the ability to collect high-throughput data in consistent assays.

The shared ancestry of organisms forms the basis of many ways we extend results from one system to another. Across species, this shared ancestry is the basis for functional annotation using homology^6^; within species, it is the basis for a shared reference to align functional genomics data^1,7^. Both of these foundational ideas exploit the shared existence of the same set of genes across systems, placing data collected from heterogeneous sources into a common framework. Whenever a gene is described as linked to a disease^5,8^, annotated with a Gene Ontology (GO) function^9,10^, or described with respect to structure or biochemical activity^11,12^, we imply a standardized description of the “same” gene found in different systems. Analytically, this frequently creates an oddity within machine learning of gene function: we are often learning over the same sample space, again and again (say, human genes), extending an initial positive set to include more and more of what were originally negatives^5^. This is unlike supervised learning in any other field where the intent is to learn a classifier that can be applied to “new” samples (as opposed to the same genes/samples over again). As a result, generalizability can only be assessed across systems (such as conditions) rather than samples; i.e., we ask does this new experiment (feature data for a gene) also imply it possesses a given function? Combined with using primarily sparse positive annotations without explicit negatives^13–15^, this separate “closed universe” problem of resampling across novel feature spaces makes it difficult to interpret annotation performance from traditional machine learning metrics alone.

A second challenge (and opportunity) relates to the magnitude of genome-scale data. In modern genomics, many assays are designed to be comprehensive across the genome, with significance arising from the combination of information across genes. This is used in differential expression^16^, enrichment analysis^17,18^, and more generally, network analyses that aim to capture gene associations of all types^1,19^. Thus, networks can be interrogated for overlaps in disease genes or other sets, with even a small number of genes contributing to generating a significant result if they are “surprisingly” close in the network. More broadly, there are two potentially complementary models for gene associations: in the first model, functions and phenotypes are well captured by a small set of genes (Mendelian diseases or large effect loci in GWAS^20,21^), while in the second model functions are distributed over a large set of genes (polygenic model^21,22^, omnigenic model^23^). In both models, proteins frequently participate in multiple functions, resulting in overlap between gene sets^24,25^, reflecting poor human definitions for functions or true multifunctionality. Likewise, diffuse interactions may reflect noisy data or true omnigenic robustness^26^. To understand these questions about the discreteness and extent of gene function, we need a framework that lets us interpret conclusions drawn in one context jointly with others.

In this article, we assess the generalizability of gene associations based on the graphical properties of performance curves. We find that genomic ROC curves endemically produce highly significant straight line segments which define gene set re-organizations. Each straight segment groups together annotated and unannotated genes that are equally likely to have the investigated function. By extracting these straight lines using a normalized Kolmogorov-Smirnov statistic, we can rapidly evaluate the generalizability of gene sets to other contexts across virtually any study. We find that straight lines are pervasive across the data sources investigated (>90% in all functional learning tasks) and suggest the existence of large gene modules (up to 40% of the genome). Finally, we confirm our findings by extracting ROC curves directly from figures in the published literature. We find interpretively important straight lines in 71/77 published performance curves covering a wide body of methods and data. Together, these results and methods for the interpretation of performance curves extend our ability to rapidly and visually probe gene set generalizability across studies and systems.

## Results

To illustrate how the shape of the ROC curve informs about the structure of the data, we first consider a toy example of protein function prediction (Fig. 1). A machine learning classifier is applied on a high-throughput dataset measuring the likelihood that two proteins interact (Fig. 1a). To make predictions, the classifier relies on annotations obtained by a low-throughput assay that labeled a subset of genes with 50% True Positive Rate (TPR) and 10% False Negative Rate (FNR)(Fig. 1a). The classifier correctly identifies that there is a functional module: the highest prediction scores contain an even mix of correctly annotated genes and unannotated functional genes, while lower scores contain an even mix of incorrectly annotated and unannotated non-functional genes.

**Figure 1.**
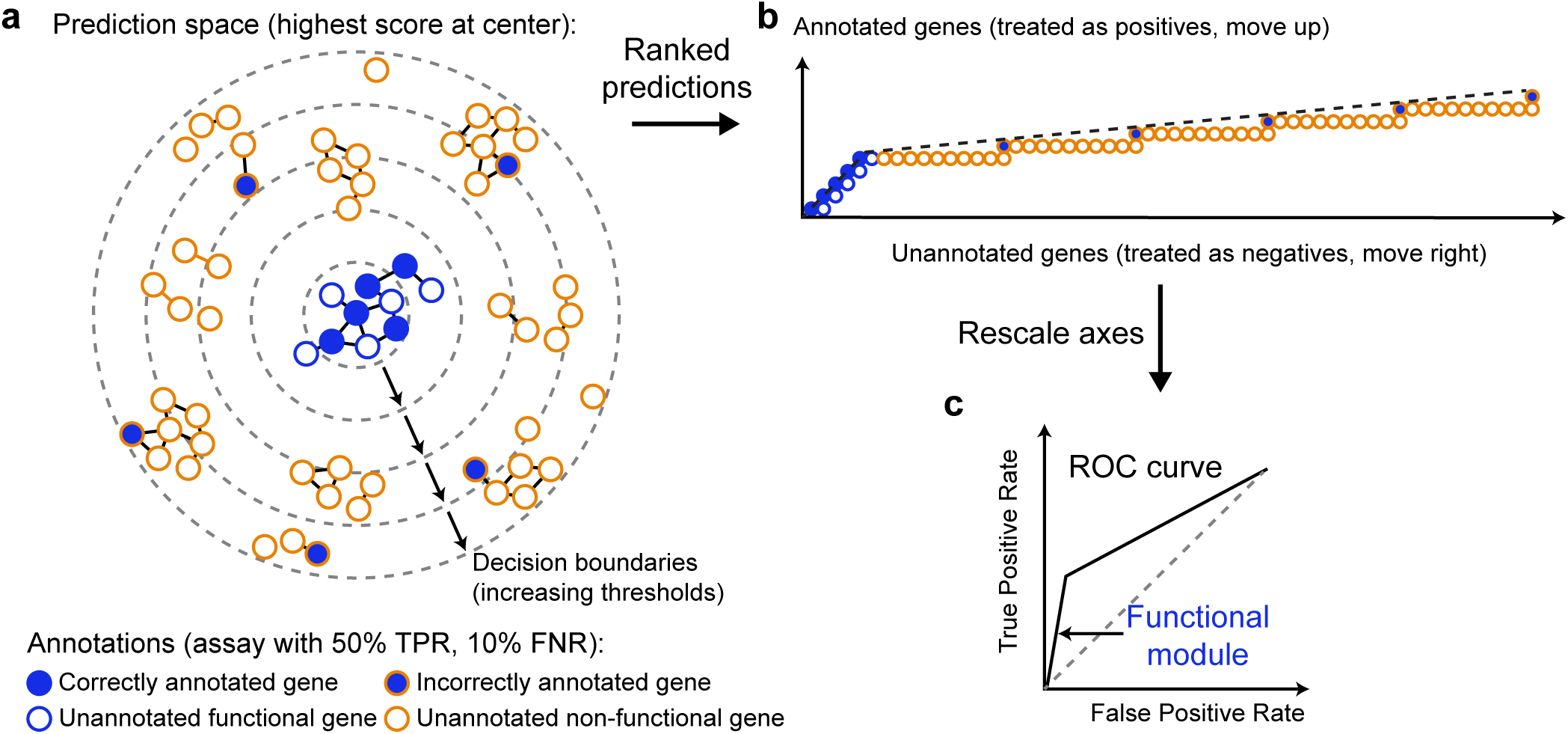
Straight lines in ROC space reveal the presence of functional modules. **a** Schematic representation of a gene function prediction task. Nodes represent genes, edges represent the strength of interactions between genes as determined by a high-throughput assay (for simplicity, only strong interactions are shown). The fill color of nodes shows the genes’ current annotation (as established by a low-throughput assay), the outline color shows the true status. Genes are organized according to prediction scores from a machine learning classifier, with genes most likely to be functional (highest predictions scores) at the center. **b** Taking current annotations as the ground truth (positive=annotated, negative=unannotated, closed-world assumption), predictions can be summarized as an ROC curve. The ROC curve can be conceptualized as a walk in (FPR, TPR) space: starting from the gene with the highest prediction score, the curve moves up every time an annotated gene is encountered, and right every time an unannotated gene is encountered. **c** The presence of a functional module that mixes annotated and previously unannotated gene is revealed by the presence of straight lines in the ROC curve.

To build an ROC curve, we apply the closed world assumption^14^: every unannotated gene is considered to be a negative. The curve can be obtained by ranking all genes by prediction score, moving up when encountering a positive (annotated gene) and right when encountering a negative (unannotated gene)(Fig. 1b), then rescaling axes to range between 0 and 1 (Fig. 1c). The presence of a functional module in the data is immediately visible on the ROC curve because of the presence of straight lines. The initial straight line on the ROC curve suggests that a subset of annotated genes shows high evidence of interaction, but also that the annotation can be “naturally” extended to some unannotated genes as, locally, annotated and unannotated genes are evenly mixed.

### Straight lines in ROC space define discrete classes of genes with similar functional properties

In an effort to identify robust functional gene sets, we start our investigation with a focus on cell-type-defining genes. Recent cell type atlasing efforts based on single-cell RNA sequencing (scRNAseq) across tissues and organisms suggest that, in their mature state, most cell types act as remarkably discrete transcriptomic entities and constitute well defined and conserved building blocks of biology^27,28^. In transcriptomic space, this discrete nature translates into well-separated clusters and cell-type-specific marker genes. Marker genes such as Slc17a7 (glutamatergic neuron marker) or Gad1 (GABAergic neuron marker) have distinctive binary expression patterns where the gene is only expressed in the cell type of interest, but none of the background cells^29^. Overall, these patterns suggest the existence of discrete gene modules associated with cell types. Ideally, such marker modules would be perfectly replicable across tissues, individuals, and technologies.

To evaluate the replicability of marker modules, we extracted markers from the Tabula Muris atlas^30^. The Tabula Muris provides the perfect opportunity to study marker gene generalizability, as it contains 100,605 cells sampled from 7 mice (3 males and 4 females). Furthermore, the atlas sampled cells from 20 organs and sequenced 55,656 cells using the 10X technology and 44,949 cells using the Smart-Seq technology. In the following, we focused on the “B-cell” cell type, because this cell type was detected in 42 combinations of individuals (7 individuals), tissues (7 organs) and sequencing technologies (10X and Smart-Seq) for a total of 10,323 cells.

First, we extracted the top 20 cell type markers (see Methods) from the “3_10_M” individual in the “Fat” tissue, sequenced using the Smart-Seq technology. This corresponds to a typical marker gene extraction scenario, in which a study relies on a single tissue and sequencing technology. To study the generalizability of these 20 markers, we asked whether they are also predicted as top markers in the remaining data. We generated one ROC curve per individual, tissue, and technology combination that contained more than 20 cells (25 combinations). In each combination, we treated the top 20 fat markers as positives and all other genes as negatives; we used the effect size of the ROC test (commonly used to compute markers in Seurat^31^ or LIGER^32^) to rank genes.

Marker replicability AUROCs ranged from 0.83 to 1 (median 0.97), suggesting high replicability across tissues, individuals, and technologies. Performance differences were mostly explained by variability across tissues (61% variance explained), then sequencing technologies (15% variance explained), and individuals (6% variance explained). The marker set was perfectly replicable within fat (AUROC∼1 across all individuals), but suboptimal in other tissues (AUROC<1).

Remarkably, we noted that the initial part of each suboptimal ROC curve was composed of straight lines (Fig. 2a). In essence, this indicates sets of genes whose order is irrelevant to the final performance statistics; the functional validation “sees” this set of genes as a uniform set. We extracted straight lines from the “Lung” tissue using the normalized Kolmogorov-Smirnov statistic (see Methods). In the initial portion of the ROC curves, we identified two straight lines spanning approximately 5% of the genome (Fig. 2b). The first line (perfect straight line, KS=NA) highlighted that approximately 50% (∼10/20) markers picked in fat were perfectly replicable in lung (Fig. 2b). This line went straight up along the y-axis, suggesting that none of the negative genes are equivalently good markers in the lung. We call these genes primary markers.

**Figure 2.**
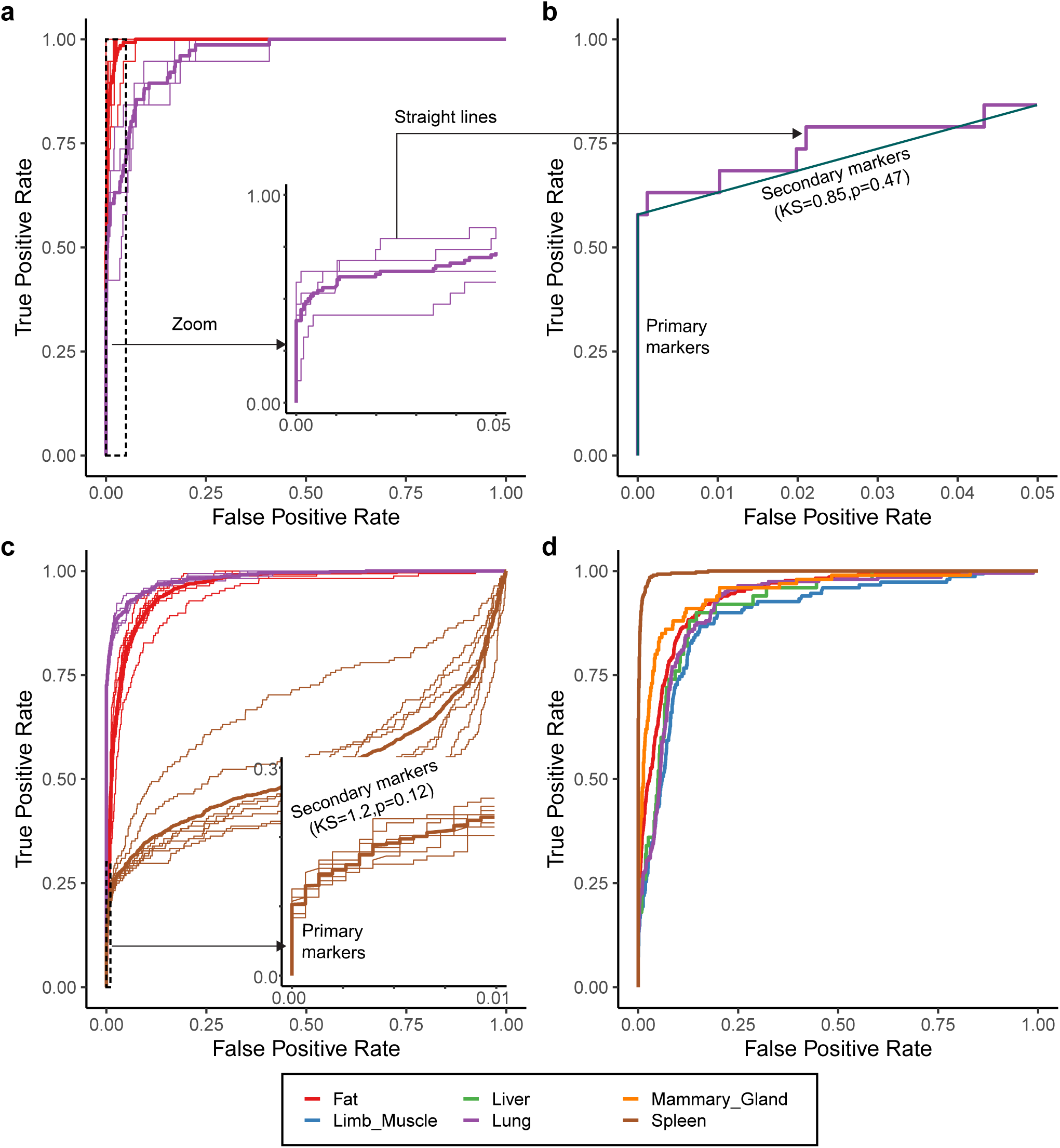
Straight lines identify generalizable markers and tissue-specific markers. **a** Evaluation of the generalizability of the top 20 Fat markers obtained from a single individual. Thin lines show individual ROC curves (one mouse, tissue, technology combination), thick lines show average ROC curves per tissue. A high AUROC indicates that markers generalize well, i.e., they are also top markers in other tissue x technology combinations. Inset: a zoom on the ROC curve up to 5% FPR highlights the presence of straight lines in the Lung. **b** The presence of straight lines can be assessed using the Kolmogorov-Smirnov (KS) test. The initial straight lines can be used to identify primary markers (perfect markers shared across tissues) and secondary markers (tissue-specific markers). Secondary markers contain a mix of previously annotated genes and unannotated genes with equal marker strength. **c** Same assessment as **a**, but taking primary and secondary markers from the Lung as the reference marker set. Inset: a zoom on the ROC curve up to 1% FPR highlights the presence of straight lines in the Spleen. **d** Same assessment as **a**, but taking primary and secondary markers from the Spleen as the reference set.

In contrast, the second line (KS=0.85,p=0.47,n=5) contained approximately 25% of the original markers, but also 5% of the negative genes. The presence of a straight line that contains both positive and negative genes suggests that all genes “contained” in the straight line have equivalent marker strength. From a biological point of view, this line suggests the existence of context-dependent markers: the list of markers extracted from fat can be completed with previously unidentified genes that could be used interchangeably as secondary markers in the lung. Accordingly, lung-specific secondary markers were consistent across all individuals and technologies (Sup. Fig. 1a).

Next, in an attempt to create a larger and more robust set of markers, we refined the initial marker set (top 20□markers from fat) using the secondary markers identified in lung. We picked all genes contained in the first two straight lines, representing a new candidate marker set with 480□genes (16/20 initial genes, 464 additional genes). This larger marker set proved to be highly replicable across most tissues (AUROC>0.96, ΔAUROC=-0.03-0.03), with the notable exception of the mammary gland (ΔAUROC=-0.09) and the spleen (ΔAUROC=-0.36, Sup. Fig. 1b). This pattern highlights the context dependency of secondary markers: they are shared for fat, the lung, limb muscles, and the liver, but not with the mammary gland or the spleen.

Having established a strong set of primary and secondary markers for 4 out of 6 tissues, we looked for evidence of spleen-specific secondary markers. Using ROC curves, we asked which unannotated genes had equivalent performance to the 480 markers extracted from the lung. Again, the initial segments of the spleen ROC curves were almost perfectly linear (KS=1.2, p=0.12, n=18 for the average ROC curve, Fig. 2c). We extracted the first two straight lines, which contained ∼25% of the lung marker set (∼10% as primary markers and ∼15% as secondary markers) and 1% negative genes with equivalent performance in the spleen, representing a total of 216 genes. This new marker set resulted in almost perfect performance in Spleen (0.99) and lower performance in other tissues, individuals and technologies (AUROC range=0.89-0.95, Fig. 2d, Sup. Fig. 1b), suggesting that most of the newly identified markers are spleen-specific.

This example shows how straight lines in ROC space enable one to quickly assess marker generalizability. They decompose a candidate gene set into discrete classes of genes with respect to a given functional property (here, cell-type-specificity). A simple look at a set of ROC curves suggests the existence of shared primary markers and secondary tissue-specific markers. The size of straight lines can be directly interpreted: there are around 10 to 50 primary markers and 100 to 500 secondary markers.

### Functional Equivalence Classes (FECs) are pervasive across the functional landscape

For cell-type-associated genes, the discrete nature of cell types translated into the existence of discrete sets of genes. But the discrete nature of the functional property assayed may have contributed to the clarity of the discrete classes. An alternative would be to take a network approach, capturing many conditions simultaneously. Network biology identifies fundamental functional building blocks by analyzing the global topology of molecular interaction networks, including gene-gene or protein-protein interactions^19^. The central hypothesis is that there are robust building blocks whose interactions are shaped by evolution. This hypothesis serves as the foundation of widespread applications such as gene set enrichment analyses^17,18^, which look for functional enrichment across a pre-defined hierarchy of discrete gene sets (such as the Gene Ontology^9,10^ or MSigDB^17^). However, it remains difficult to test how well discrete gene sets are supported by the data, and how context-dependent they are.

As we have seen previously, straight lines in ROC space define discrete classes of functionally equivalent genes. Usually, perfect straight lines arise when predictions are tied. In contrast, here we are interested in straight lines in the absence of ties, which arise when the score distribution of a subset of negatives and positives is identical. Mathematically, these lines are conserved under class label permutation. Figure 3a shows an example of a performance curve for a gene function prediction task for the “growth factor activity” function (GO:0008083, Molecular Function ontology). Strikingly, this ROC curve is composed of multiple straight lines. Each straight line can be interpreted as a pool containing positive and negative genes with equal strength, i.e. the class labels are locally interchangeable (Fig. 3b). If we permute the class labels globally, the ROC curve becomes a random walk along the diagonal (AUROC ∼ 0.5, Sup. Fig. 2a). However, if we only permute labels from genes belonging to a straight line, the ROC curve is essentially unchanged (Fig. 3c, Sup. Fig. 2b). Permutations can be seen as a local random walk, allowing to automatically detect straight lines using the normalized Kolmogorov-Smirnov test statistic (see Methods). In the following, we refer to straight lines in ROC space as Functional Equivalence Classes (FECs).

**Figure 3.**
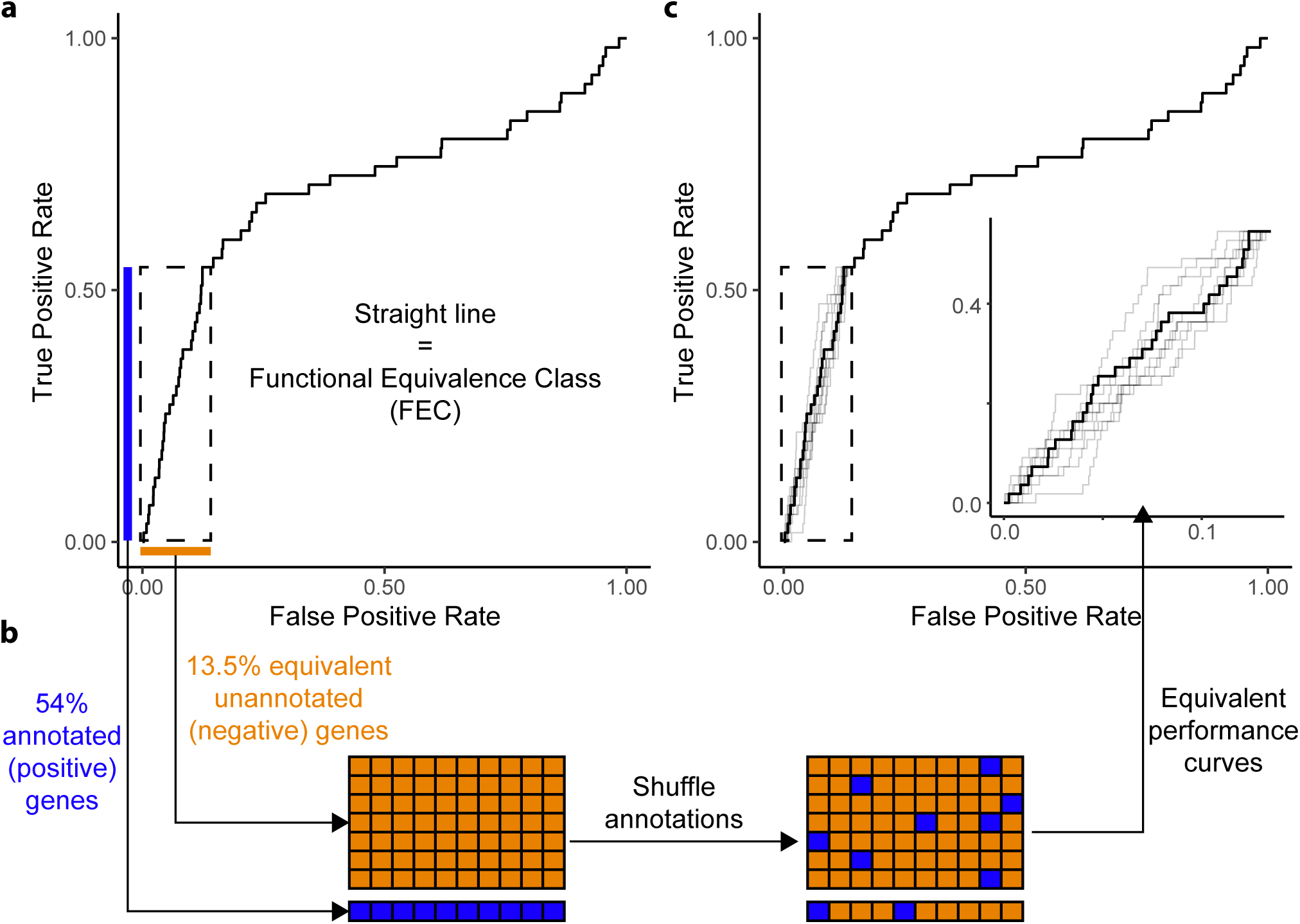
Straight lines in ROC space are Functional Equivalence Classes (FECs). **a** Example of ROC curve obtained from a function prediction task. The initial part of the ROC curve is a FEC, a straight line containing a mix of 54% previously annotated genes and 13.5% previously unannotated genes. **b** The presence of a straight line indicates that the classifier sees positives and negatives as functionally equivalent, as if they originated from a single class. Formally, the presence of a FEC can be tested by shuffling annotations. **c** Under local annotation permutation (within the FEC), the ROC curve is essentially unchanged. The thick black line shows the original ROC curve, thin gray lines show 10 simulations of annotation permutation. The inset shows a zoom of the FEC.

We now turn to a broad set of functions as defined by the Gene Ontology (GO), and investigate the presence of FECs across two types of network data offering wide meta-analytic resources and capturing different aspects of function: Protein-Protein Interaction (PPI) networks and co-expression networks. PPI networks are binary networks where nodes are proteins and edges connect pairs of proteins that physically interact. Data used to build PPI networks include yeast two-hybrid assays (assessing whether a pair of proteins interact) and affinity capture methods (finding all proteins that interact with a given protein). In contrast, co-expression networks are weighted networks where nodes are genes and edges reflect the propensity of two genes to be expressed in the same contexts (conditions, tissues or cell types). Co-expression networks rely on genome-wide assessments, usually from microarray or RNAseq experiments.

We built a PPI network by aggregating all interactions from the BIOGRID^33,34^ database annotated as “Mouse” and “Physical Interaction”, resulting in a network containing 10,172 proteins and 57,337 interactions. As PPI networks are typically sparse, we used a propagation algorithm to obtain a dense network, which accounts for indirect interactions between proteins (see Methods). We downloaded the mouse co-expression network from the CoCoCoNet^35^ database. The network was obtained by aggregating 3,359 samples over 85 experiments, resulting in a dense network containing 17,834 genes. To allow comparisons between the two modalities, we restricted the two networks to 9,058 common genes. We restricted our study to 4,238 well-powered GO terms containing at least 20 genes.

To assess whether a function is supported by a network’s topology, we use the guilt-by-association framework implemented by the EGAD^36^ algorithm. Briefly, EGAD uses a neighbor voting algorithm to assess whether genes that are annotated with the same function tend to be neighbors in the network (Fig. 4a). Some of the annotated genes are held-out and serve as positives, while all other genes are annotated as negatives (closed world assumption). Taking neighbor votes as a predictor for held-out genes, we build one ROC curve for each function and network. A high AUROC indicates that the function is supported by the network, i.e. genes with this function tend to be highly modular. Overall, GO functions were strongly modular in both the PPI (median AUROC=0.72) and co-expression networks (median AUROC=0.70, Fig. 4b). Performance was only partially correlated (rho=0.35, Sup. Fig. 3a), consistent with the fact that PPI and co-expression capture different aspects of function.

**Figure 4.**
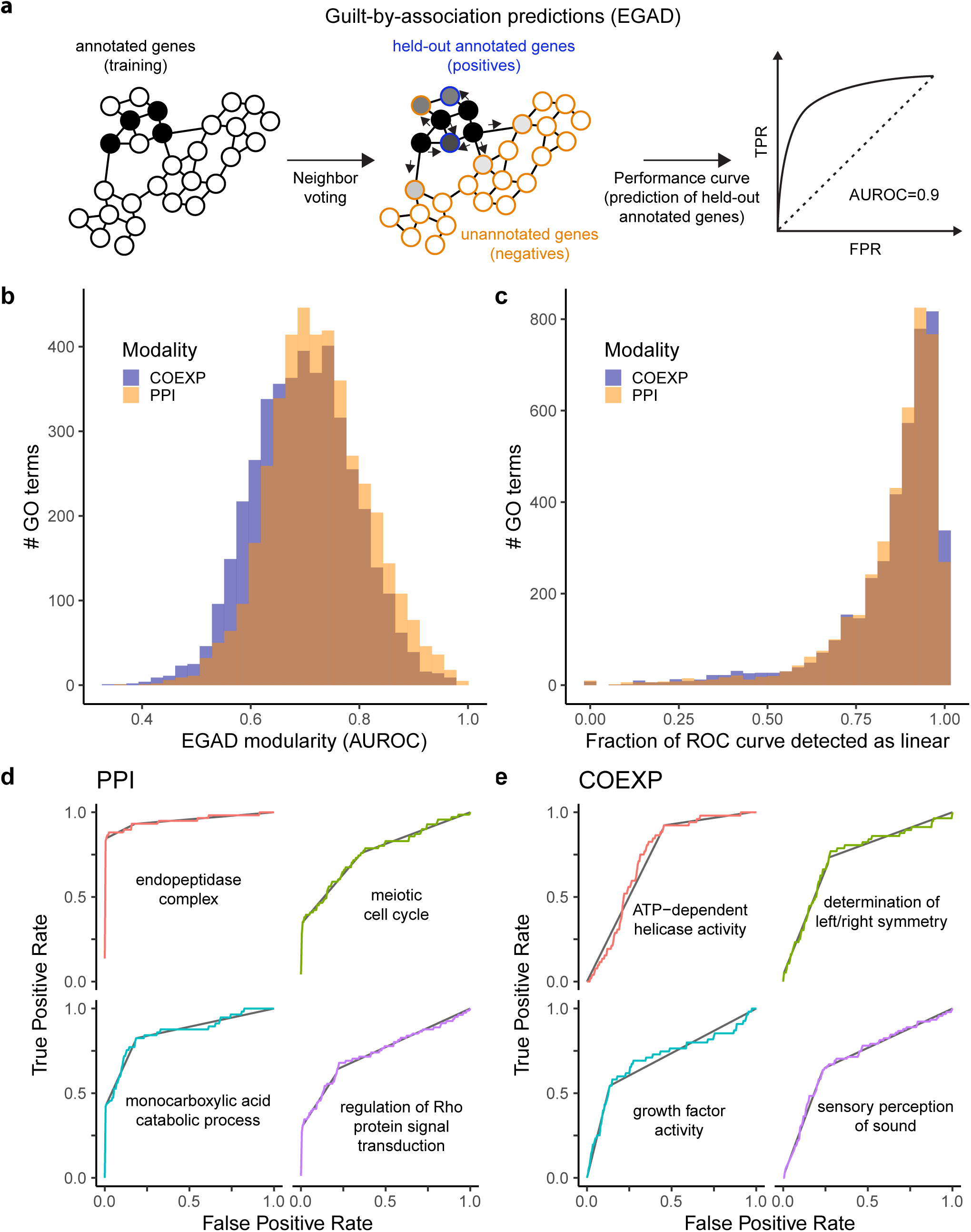
FECs are pervasive across the functional landscape. **a** Schematic of the function prediction task. EGAD predicts functional annotations based on the connectivity of genes in network data (neighbor voting algorithm). Using held-out annotated genes as positives, performance can be summarized as an ROC curve, which reflects the degree of modularity of a functional gene set. **b** Degree of modularity (EGAD AUROC) of functional gene sets defined by the Gene Ontology (GO) in meta-analytic co-expression (COEXP) and protein-protein interaction (PPI) data. **c** Fraction of ROC curves detected to be straight lines. **d**,**e** Examples of ROC curves composed almost exclusively of straight lines. Each facet shows a specific GO term, colored curves show the ROC curve for this term, black lines show the FECs detected using the KS test.

Unexpectedly, FECs that spanned at least 5% of the genome were detected in 99.8% functions (undetected in 10/4238 PPI curves, 8/4238 co-expression curves) and spanned 85% of the genome on average (Fig. 4c). 95/8478 (1.1%) functions were even detected to be entirely composed of straight lines, such as “meiotic cell cycle” (2 FECs, Fig. 4d) or “determination of left/right symmetry” (2 FECs, Fig. 4e).

3980/8476 functions (46% in co-expression, 48% in PPI) contained exactly two FECs (Fig. 5a). The length of individual FEC varied substantially across functions, and had a clear bimodal shape in both modalities (Fig. 5b). The first mode contained 62% of FECs and spanned 5% to 40% of the genome. Upon further investigation, this mode roughly corresponded to the length of the primary FEC of each curve (FEC containing high-ranking genes, i.e. most likely to have the function)(Sup. Fig. 4). Smaller primary FECs were found more frequently: 50% FECs spanned <13% of the genome, 80% FECs spanned <23%.

**Figure 5.**
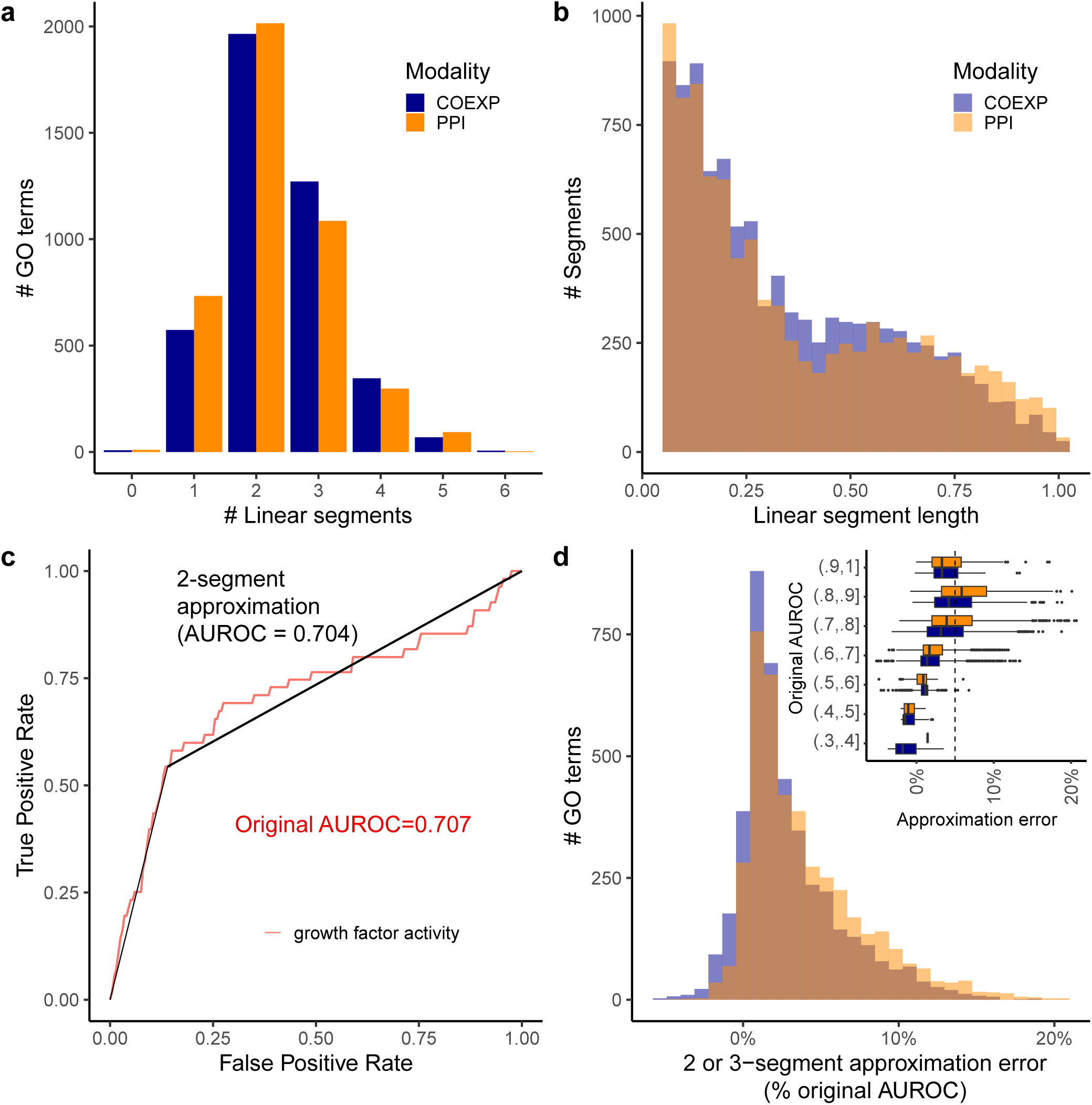
FECs offer a data-driven view of the extent of biological functions. **a** Distribution of the number of FECs detected in ROC curves corresponding to each GO term. **b** Distribution of length of individual FECs, as measured by the fraction of the x-axis (FPR axis) spanned by the FEC. **c** Example of 2-segment approximation for the “growth factor activity” GO term. The initial ROC curve and AUROC are shown in red, the straight line approximation and approximated AUROC are shown in black. **d** Distribution of approximation error on the AUROC when swapping ROC curves by their 2 or 3-segment approximation. A low approximation error suggests that performance is driven by the presence of 2 or 3 discrete modules in the data. The inset further breaks down the approximation error by stratifying on the original AUROC.

Because FECs often spanned large portions of the genome and most curves contained exactly 2 or 3 FECs, we wondered how many functions could be explained by the presence of two or three discrete classes of genes. We identified the start and end of the longest FEC in each curve, replaced it by a straight line, then connected this line to the (0,0) and (1,1) points using straight segments. In the case where the FEC already contains the (0,0) or (1,1) point, the ROC curve can be correctly approximated by two straight lines (function-associated vs non-function associated, Fig. 5c). We found that the two or three line-approximation worked to a surprising degree: 74% of curves (50% for high AUROC curves) could be approximated with a relative error on the AUROC lower than 5% (Fig. 5d).

Despite partially uncorrelated performance, the presence and size of FECs was remarkably consistent across the PPI and co-expression modalities. These results suggest that modular structure is widespread in the data, although modules only partially overlap with existing annotations. For most functions, the data even suggest a binary partition of genes, with function-associated genes constituting up to 40% of the genome.

### FECs are pervasive in the published literature

Our previous experiments showed that FECs are detected across a large fraction of biological functions. We next wondered whether FECs would hold across an even larger body of methods and data, and looked for evidence of straight lines in individual studies.

To assess the presence of FECs in published work, we extracted ROC curves from 50 research articles, composed of an unbiased selection of 35 articles from the PLoS One journal and 15 manually curated high-profile articles (Sup. Note). In total, we extracted 77 ROC curves from main and supplementary figure panels using the Engauge Digitizer^37^ software (Fig. 6a). As in previous sections, we assessed the presence of straight lines using the normalized KS statistic (Methods).

**Figure 6.**
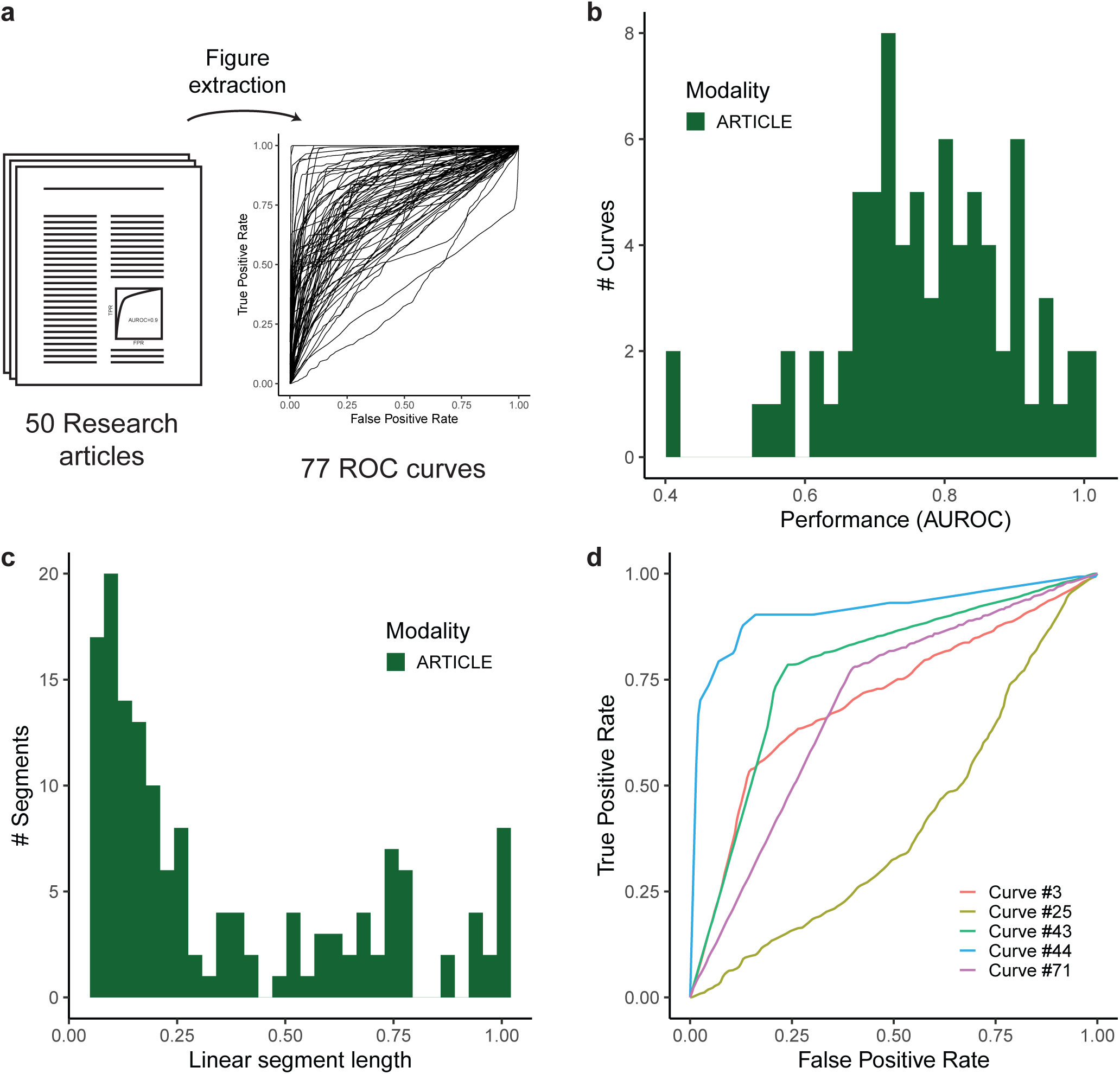
FECs are pervasive in research articles. **a** Schematic of ROC curve extraction using the Engauge Digitizer tool. ROC curves were extracted from 15□selected publications and 35 publications published during one calendar year in genomics-related subject areas of the PLoS One journal. **b** Distribution of estimated AUROCs from curves extracted from the literature. **c** Fraction of ROC curves detected to be straight lines. **d** Examples of ROC curves composed almost exclusively of straight lines.

While the median performance reported in published studies (median AUROC=0.78, Fig. 6b) was slightly higher than the median performance in PPI and co-expression data (median AUROC=0.71), the presence of FECs was equally widespread. 92% (71/77) curves contained FECs, spanning 71% of the curve on average (Sup. Fig. 5a).

Remarkably, the shape of ROC curves had similar properties to curves obtained from PPI and co-expression data. 31/77 curves were composed almost entirely of straight lines (covering >90% of the curve), with 39/77 curves contained exactly 2 or 3 segments (Sup. Fig. 5b). Even the distribution of individual FEC length was remarkably similar to the PPI and co-expression distributions, with primary FECs spanning approximately 5 to 40% of the genome (Fig. 6c, Sup. Fig. 5c), and 68% of curves correctly approximated by two or three linear segments (Fig. 6d, Sup. Fig. 5d). These results suggest that FECs are an omnipresent characteristic of genomics data, found across data sources and machine learning methods.

Interestingly, ROC curves extracted from the published literature contained another striking pattern: for 4/71 (6%) curves, flipping a segment of the curve significantly increased the AUROC performance (Fig. 7a). In the prediction space, a flip corresponds to inverting the ranking of genes contained in the segment: genes that were predicted as most likely to have the function are now predicted as least likely. In some instances, optimal performance was achieved by flipping the whole curve (i.e., completely inverting predictions); in others, the curve was locally S-shaped and the optimal flip only contained around 50% of the curve (Fig. 7b). The latter case suggests that the best predictions are located in the initial and final part of the S, and would typically arise by mistakenly treating a two-sided assessment (where both highly positive and highly negative predictions should be considered “high”) as one-sided (where only positive values are considered high). “Flippable segments” were notably absent in ROC curves extracted from co-expression and PPI data, suggesting that they are likely related to extreme data distributions or methodological issues.

**Figure 7.**
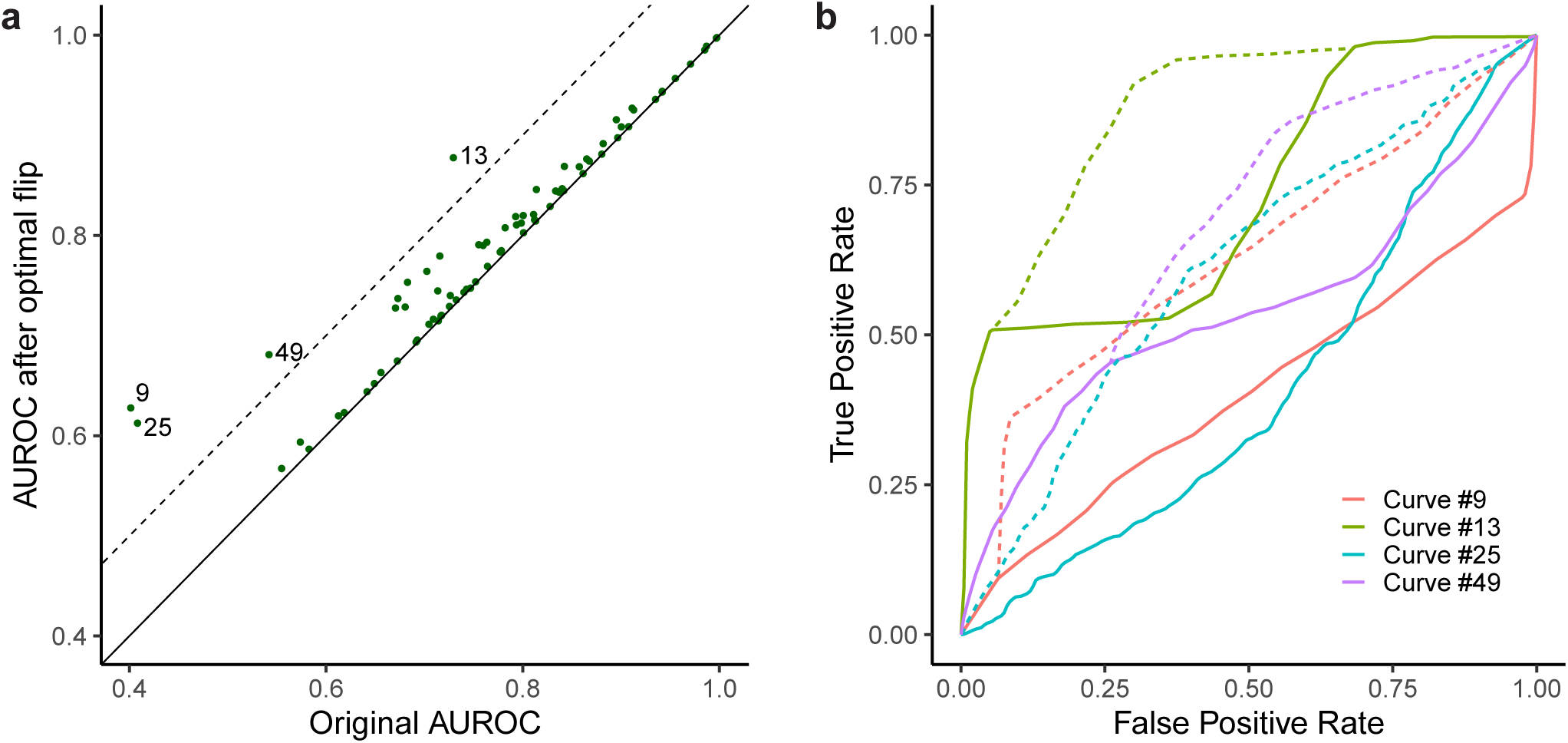
Flips in ROC curves identify sections where predictions can be inverted. **a** Scatter plot showing optimal AUROC increase obtained by flipping a single ROC segment. The dashed line corresponds to a 0.1 AUROC increase. **b** ROC curves of the 4 curves with AUROC increase > 0.1 highlighted in **a**. Solid lines show the original ROC curve, dashed lines show the ROC curve after flipping the optimal ROC segment. In prediction space, these flips correspond to inverting the order of predictions for the genes contained in the ROC segment.

## Discussion

In this study, we showed that the shape of ROC curves offers a visual and data-driven interpretation of the extent of biological functions. The presence of straight lines in the ROC curve suggests that the data are compatible with the extension of a functional gene set to unannotated genes. For example, starting with a handful of marker genes for a given cell type, straight lines let us rapidly identify unannotated genes of equivalent strength. We call these straight lines Functional Equivalence Classes (FECs), because they define discrete classes of genes that are equivalent with respect to the functional property investigated.

Our examples show that the extensibility of gene sets is context specific: B-cell markers that worked well in the lung did not necessarily work in the spleen. Instead, we found that a subset of primary markers was conserved across tissues, while secondary markers varied from tissue to tissue. One of the strengths of FECs is that the generalizability and extensibility of a gene set can be probed with one look at the ROC curve. Either the gene set works perfectly well in the new context (AUROC=1), or performance is suboptimal and FECs suggest how the gene set can be reorganized in the new context. FECs can be automatically extracted using the normalized Kolomogorov-Smirnov statistic, suggesting which genes to remove from the initial annotation (not generalizable) and which genes to add (equivalent in the current context).

Straight lines were equally prevalent in ROC curves obtained by testing marker consistency in single-cell data, ROC curves mined from published articles sampling across various sources of data and methods, and ROC curves obtained by testing the modularity of GO sets in meta-analytical PPI and co-expression data. The existence of straight lines is thus unlikely to be an artefact of specific data or methods. At least 92% of ROC curves contained straight lines, and curves were often piecewise linear. Overall, the data support a view where most functions map to discrete representations in the genome, with the initial FEC representing a pool of genes primarily engaged in the function (primary FEC).

The omnipresence of FECs suggests a functional space where functions are distributed over discrete gene sets. This view is compatible with the discrete organization of genes in gene sets (such as GO sets, MSigDB signatures, or marker sets) and reminiscent of the polygenic model, where disease risk is distributed over a larger set of genomic loci^22,38,39^. However, these discrete sets are observed in one biological (e.g., a given tissue in the marker space) or technological (PPI, co-expression data) context. The marker example suggests that, while primary and secondary markers formed discrete sets of genes, the secondary marker sets varied from one tissue to the other. Integrated across enough contexts, the degree of functionality of genes may start to appear continuous, consistent with the omnigenic model, which posits that all expressed genes are likely to contribute a disease due to the interconnections of regulatory networks^23^.

ROC curves have a rich history in genomic assessments, and are frequently interpreted through the lens of AUROCs. However, AUROCs are often deemed unintuitive in the presence of extreme class imbalance, which led to a more particular focus on the evaluation of top predictions, either through partial ROC curves (ROC50^40^, pAUC^41,42^) or precision-recall curves^43,44^. Interestingly, what is seen as a weakness becomes a strength when the ROC curve as a whole is considered. Indeed, for given positive and negative score distributions, the shape of the ROC curve is independent of class imbalance, facilitating visual interpretation. As seen in this article, when negatives and positives have the same score distributions locally, we obtain “null” segments in the form of straight lines that can be rapidly identified visually or statistically. Strictly speaking, FECs could also be extracted from other performance curves, such as the precision-recall curve. However, null segments are harder to assess visually, as they become curves in precision-recall space (see e.g.^45^), with highly unintuitive curvatures that depend on class imbalance^46^. The idea that the shape of the ROC curve can be visually interpreted was previously noted by Janssens and Martens, who distinguished “rounded” and “non-rounded” ROC curves, attributing the occurrence of “angles” to the presence of a dominant binary predictor^47^. In this study, we find that the presence of such “angles” is widespread in genomic data, and usually accompanied by the presence of straight lines, suggesting an underlying modular organization of the data.

FECs define a formal framework to visualize and probe the context-specificity of functional gene sets. They are simple to visualize and extract, providing a novel way to summarize complex data. They are widely applicable, as ROC curves are frequently used in genomic assessments, paving the way for comparative and meta-analytic studies. Applied across a range of contexts, they provide a first step towards teasing out shared and context-specific gene set components.

## Methods

### Datasets

We downloaded the Tabula Muris^30^ single-cell RNA sequencing (scRNAseq) dataset from FigShare, specifically Version 2 of the 10X^48^ and Smart-Seq2^49^ data, along with metadata and annotations. The expression data and annotations from individual tissues were then merged into two SingleCellExperiment objects (one for 10X and one for Smart-Seq2) for downstream analyses in R, keeping all cells for which an annotation was available (100,605 cells). For the marker analysis, we applied CP10K (counts per 10k) normalization for the 10X data and CPM (counts per million) normalization for the SmartSeq data using custom code.

We downloaded the mouse PPI data BIOGRID-ALL version 4.4.197 from the BIOGRID□website^33,34^. We filtered the BIOGRID data for mouse (taxonomy ID 10090), physical interaction (“Experimental.System.Type” == “physical”), then converted the data into a sparse matrix format using the “sparseMatrix” function in R. This initial network contained 57,337 interactions across 10,172 genes. To take into account indirect connections^50^, we propagated the existing interactions, setting the weight for each pair of proteins as 1/shortest path between the two proteins. To do this, we imported the sparse matrix in Python, then computed the shortest paths between any two proteins using the scipy.sparse.cs.graph.shortest_path function from the scipy package in Python.

We downloaded the mouse co-expression network from the CoCoCoNet^35^ website (last updated on 04/20/2021). We then converted Ensembl identifiers gene into symbols using the mapIds function from the AnnotationDbi R package and the org.Mm.eg.db R package (last updated on 04/21/2021). We only kept genes that had a 1:1 mapping from Ensemble to gene symbols, resulting in a dense network of 17,834 genes.

To make co-expression and PPI network performance comparable, we subset the two networks to common genes, resulting in networks of 9,058 common genes.

### Gene Ontology Annotations

We downloaded the Gene Ontology^9,10^ (GO) from the GO website (GO-basic table in OBO format, last updated 12/18/2019), then imported the ontology in R using the get_ontology function from the ontologyIndex package, with “extract_tags=“everything””. We downloaded gene ontology annotations from the MGI website (“gene-association” table, last updated 09/09/2019). Some annotations contained alternative GO IDs, which we converted to default IDs using the “alt_id” slot from the ontology. We propagated annotations to higher level terms using the “ancestors” slot. Finally, we exported the propagated annotations in sparse matrix format for downstream use in R and Python. For downstream analyses, we only kept 4,238 GO terms with ≥20 annotated genes after restricting to common genes.

### Computation of ROC curves from marker sets from scRNAseq data

We computed markers for each mouse by tissue combination independently using 1-vs-all DE (genes upregulated in one cell type compared to cells from all other cell types). Explicitly, we first subset the datasets to a given mouse using the “mouse_id” metadata, then ran the compute_markers function from the MetaMarkers^51^ package on the CP10K- or CPM-normalized Tabula Muris expression matrix, using the “cell_ontology_class” as cell type labels, and “tissue” metadata as group labels (stratifying marker search by tissue). MetaMarkers returns a ranked list of all genes, with the best markers ranked at the top. We then combined all putative markers into a single table. Because of the stratification by individual, tissue and sequencing technologies, some data combinations contained too few cells or reads for robust marker inference. We only kept markers inferred for cell types containing at least 20 cells, and genes that had a detection rate≥10% in all mice for a given tissue by cell type combination.

The first set of markers was obtained by selecting the top 20 markers for the “3_10_M” mouse from the Smart-Seq dataset, for the “B cell” cell type in the “Fat” tissue. To compute ROC curves, we asked whether these top 20 markers were also top markers in other mouse by tissue combination. For a given mouse and tissue combination, we extracted genes from the previously computed marker table, then ordered genes according to the effect size of the ROC test, “auroc” column in the MetaMarkers table. This ordered list was used as a predictor for the top 20 Fat markers; we used the “prediction” and “performance” functions from the R ROCR package to compute the ROC curve (“tpr” and “fpr” statistics) and the AUROC (“auc” statistic).

To determine the second set of markers, we visually estimated that the top 2 FECs in “Lung” spanned 5% of negatives. We extracted and pooled the top 5% markers (ranked by MetaMarkers “auroc”) in all 4 individuals (“3_39_F”, SS, “3-F-57”, 10X, “3-F-56”, 10X, “3-M-7/8”, 10X), resulting in a marker set of 480 genes. The ROC computation was similar to the prediction of the top 20 Fat markers, exchanging the initial top 20 markers with this second set of markers.

To determine the third set of markers, we visually estimated that the top 2 FECs in “Spleen” spanned 1% of negatives. We extracted and pooled the top 1% markers (ranked by MetaMarkers “auroc”) in all 8 individuals (“3-M-8”, 10X, “3-F-56”, 10X, “3_8_M”, SS, “3_9_M”, SS, “3_11_M”, SS, “3_10_M”, SS, “3_38_F”, SS, “3_39_F”, SS), resulting in a marker set of 216 genes. The ROC computation was similar to the prediction of the top 20 Fat markers, exchanging the initial top 20 markers with this third set of markers.

### Computation of ROC curves from EGAD modularity in PPI and co-expression networks

EGAD^36^ computes the modularity of a set of genes in a given network by using a neighbor voting algorithm. Since EGAD’s implementations was in R originally, we re-implemented EGAD’s modularity metric in Python. Following the original algorithm, we implemented 3-fold Cross-Validation (CV), with 2/3 of positives used for training, and 1/3 of positives held-out for testing. In detail, let *X*_*ij*_ be a positive symmetric adjacency matrix representing a weighted network, where *i* and *j* range from 1 to N (Number of genes). Let *P*_*i*_ be the one-hot encoding of the training positives. The EGAD algorithm was reproduced by computing the node degree *D*_*i*_ = ∑_*j*_ *X*_*ij*_, the neighbor votes *V* = *X.P* and finally the normalized neighbor vote 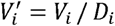 The normalized neighbor votes was used as a predictor of held-out positives, yielding one ROC curve per CV fold. For a given gene set, the final ROC curve is reported as the average ROC curve (across the 3 CV folds), the final AUROC as the average AUROC.

### Extraction of ROC curves from the literature

We systematically sampled 35 research articles from the PLoS One journal during one calendar year (genomics-related Subject Areas) and selected 15 high-profile research articles (see Supplementary Note for a detailed list of papers and figures extracted). We used the Engauge Digitizer^37^ software to extract curves from the selected figures, following the standard procedure (selection of 3 axis points for scale, automatic segment detection). In instances where the figure contained too many overlapping curves and individual curves proved too difficult to extract, we removed the figure from the analysis. After the extraction process, Engauge Digitizer generated CSV files with data points evenly distributed along the curve. To harmonize the curve resolution, we interpolated the curves such that they contained 200 total points evenly spaced along the x-axis (FPR axis).

### Assessment of linear segments using the Kolmogorov-Smirnov statistic

The one-sample Kolmogorov-Smirnov (KS) statistic computes the maximum deviation of an empirical cumulative distribution function (ECDF) from a theoretical cumulative distribution function (CDF). Conceptually, the KS statistic is related to the Brownian bridge, a random walk with fixed starting and ending points. The KS statistic measures the maximal deviation from a straight line connecting starting and ending points, with expected deviations provided by the Kolmogorov distribution. Since the ROC curve can be seen as a walk in (TPR, FPR) space (Fig. 1), the walk becomes a Brownian bridge under random labeling of positives and negatives. The “randomness” of annotation (local equivalence of positives and negatives) can thus be evaluated by computing the normalized KS statistic and comparing it to the Kolmogorov distribution.

Formally, we assessed the linearity of an ROC subcurve by rescaling it to a [0,1] by [0,1] square, then computing its deviation from the diagonal line (CDF of the uniform distribution). Mathematically, given a subcurve starting at the (*FPR*_0_,*TPR*_0_) point and ending at the (*FPR*_1_,*TPR*_1_) point, we computed the rescaled subcurve given by *FPR*′ = (*FPR* − *FPR*_0_) / (*FPR*_1_ − *FPR*_0_) and *TPR*′ = (*TPR* − *TPR*_0_) / (*TPR*_1_ − *TPR*_0_). We then computed the deviation from the diagonal (KS statistic), *D*_*n*_ = *sup*(|*TPR*′ − *FPR*′|), and the normalized KS statistic, 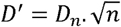. Under random annotations, the normalized KS statistic asymptotically follows the Kolmogorov distribution, i.e. the expected deviation of a Brownian bridge. The asymptotical 5% threshold to reject linearity is ∼1.36. To compute p-values, we used the C_pKS2 function used by the R function ks.test, corresponding to the one-sample test with uniform distribution and parameter “exact=FALSE”.

In the marker example, we visually estimated the range of FECs (5% for Lung, 1% for spleen), then computed the KS statistic and p-values for the extracted segments as described above. For ROC curves computed from PPI data, co-expression data and curves extracted from articles, we applied an automatic extraction procedure. We considered all possible subcurves (start/end point combinations) spanning at least 5% negatives (*FPR*_1_ − *FPR*_0_ ≥ 0.05). We then computed the normalized KS statistic *D*′ as described above. For articles, the number of positives was generally unknown and was set arbitrarily to n=100. We then tagged all subcurves with *D*′ ≤ 1 (asymptotic p-value of p≥0.27) as linear, creating a list of candidate straight lines for each ROC curve. Note that, at this stage, candidate straight lines may overlap. We established the final list of FECs using a greedy algorithm, starting by extracting the longest straight line. We then extracted the second-longest straight line that did not overlap with the previously extracted FEC, and so on, until no candidate straight line remained.

### Longest segment approximation of ROC curves

For each ROC curve, we extract the longest linear segment as described above, then reduced the ROC curve to 4 points: (0,0), (*FPR*_0_,*TPR*_0_), (*FPR*_1_,*TPR*_1_), (1,1), where (*FPR*_0_,*TPR*_0_) and (*FPR*_1_,*TPR*_1_) are the two extremities of the longest linear segment. We then computed the AUROC using the trapezoidal rule.

### Extraction of optimal ROC subcurve flip

For each ROC curve, we computed the updated AUROC after flipping each possible subcurve. Given a subcurve with extremities (*FPR*_0_,*TPR*_0_) and (*FPR*_1_,*TPR*_1_), we computed a local area under the curve (*AUC*_*local*_) using the trapezoidal rule. Specifically, we considered the area between the subcurve, the horizontal line at *TPR* = *TPR*_0_ and the vertical line at *FPR* = *FPR*_1_. Note that the complement of this area in the rectangle enclosing the subcurve is given by Δ*TPR*. Δ*FPR* − *AUC*_*local*_, where Δ*TPR* = *TPR*_1_ − *TPR*_0_, and Δ*FPR* = *FPR*_1_ − *FPR*_0_. Considering that flipping the subcurve is equivalent to rotating the rectangle enclosing the subcurve by 180°, the impact on the global AUROC can be computed as

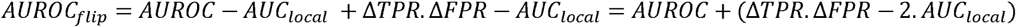

where *AUROC* is the original AUROC. After considering all start and end combinations, we identified the subcurve flip that resulted in the highest *AUROC*_*flip*_.

## Acknowledgments

JG was supported by NIH grants R01MH113005 and R01LM012736. SF was supported by NIH grant U19MH114821.

## Author contributions

SF and JG designed the experiments, performed the data analysis and wrote the paper. All authors read and approved the final manuscript.

## Supplementary Information

**Supplementary Note. List of articles and figures selected for ROC curve extraction**.

**Supplementary Figure 1. a** Upset plot showing the overlap of primary and secondary Lung markers across individuals. In the name of individuals, F/M indicates the sex, ss/10x indicates the sequencing technology (ss=SmartSeq, 10x=Chromium 10X), the remaining number indicates the age of the mouse in weeks. **b** Generalizability of the 3 marker sets (top 20 from Fat, primary and secondary markers from Lung, primary and secondary markers from Spleen) across tissues.

**Supplementary Figure 2. a** ROC curves generated by shuffling annotation labels globally. The thick black line is the original ROC curve, the thin gray line shows 10 independent simulations of label shuffling. **b** Impact of label permutation on AUROC (100 independent permutations for each case). Colors show two types of permutations: labels are shuffled globally (blue, as shown in **a**) or only within the initial FEC (red). The AUROC from the original AUROC curve is shown as a dashed line.

**Supplementary Figure 3. a** Scatter plot showing EGAD modularity for 4,238 GO terms in PPI and co-expression networks. The dashed line is the diagonal (same modularity in both networks), the correlation value is Spearman correlation. **b** Relationship between the number of linear segments (FEC) and fraction of curve detected as linear for curves computed from the co-expression (COEXP) network. **c** Same as **b** for the PPI□network.

**Supplementary Figure 4. a** Distribution of length of the initial FEC, as measured by the fraction of the x-axis (FPR axis) spanned by the FEC. Curves that did not have any FEC are omitted from the distribution. **b** For the co-expression network (COEXP), AUROCs from the original ROC curve against AUROC from the straight line approximation based on the longest FEC. **c** Same as **b** for the PPI□network.

**Supplementary Figure 5. a** Fraction of ROC curve detected as linear for the 77 ROC curves extracted from the literature. **b** Distribution of the number of FECs detected in each of the 77 ROC curves. **c** Distribution of length of the initial FEC, as measured by the fraction of the x-axis (FPR axis) spanned by the FEC. **d** Distribution of approximation error on the AUROC when swapping ROC curves by their 2 or 3-segment approximation.

