## Supplementary figures and images for "Defining the extent of gene function using ROC curvature"

### Supplementary Figures

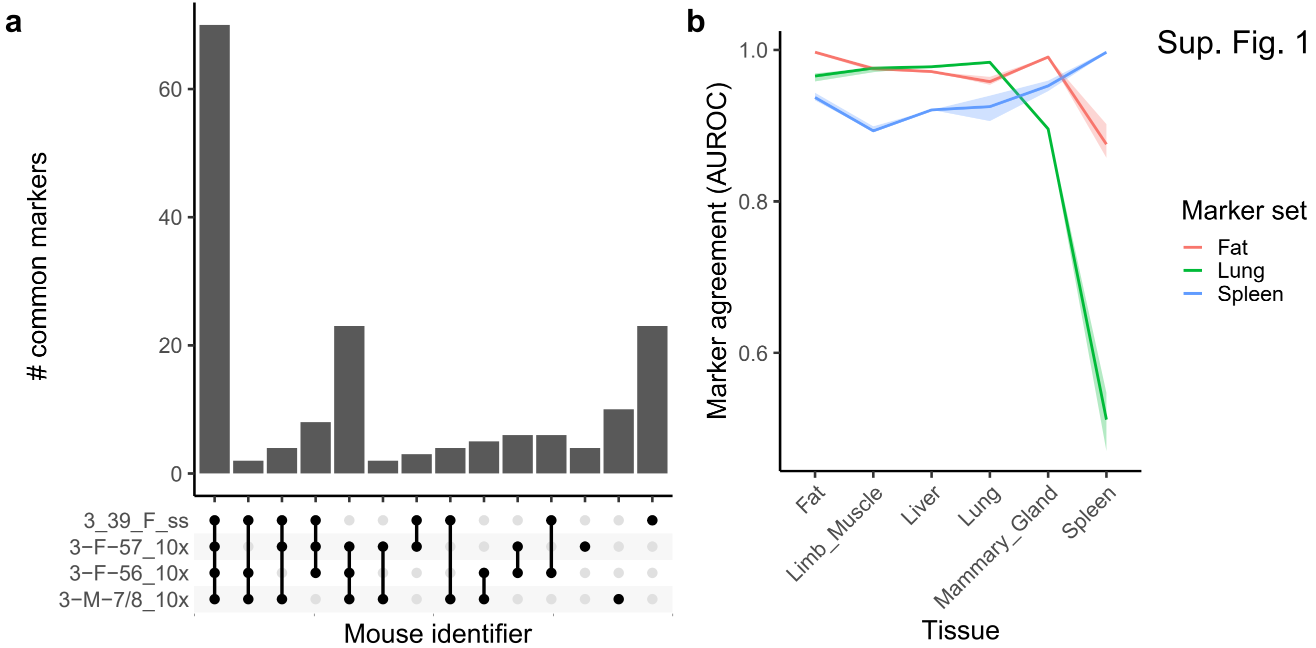

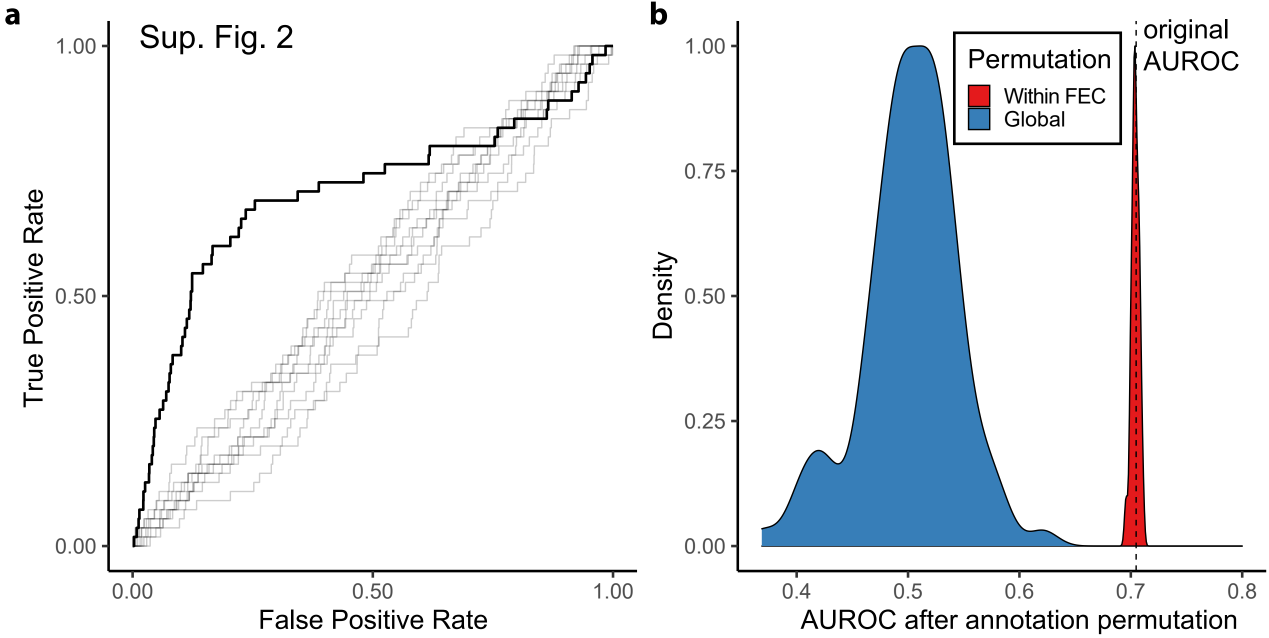

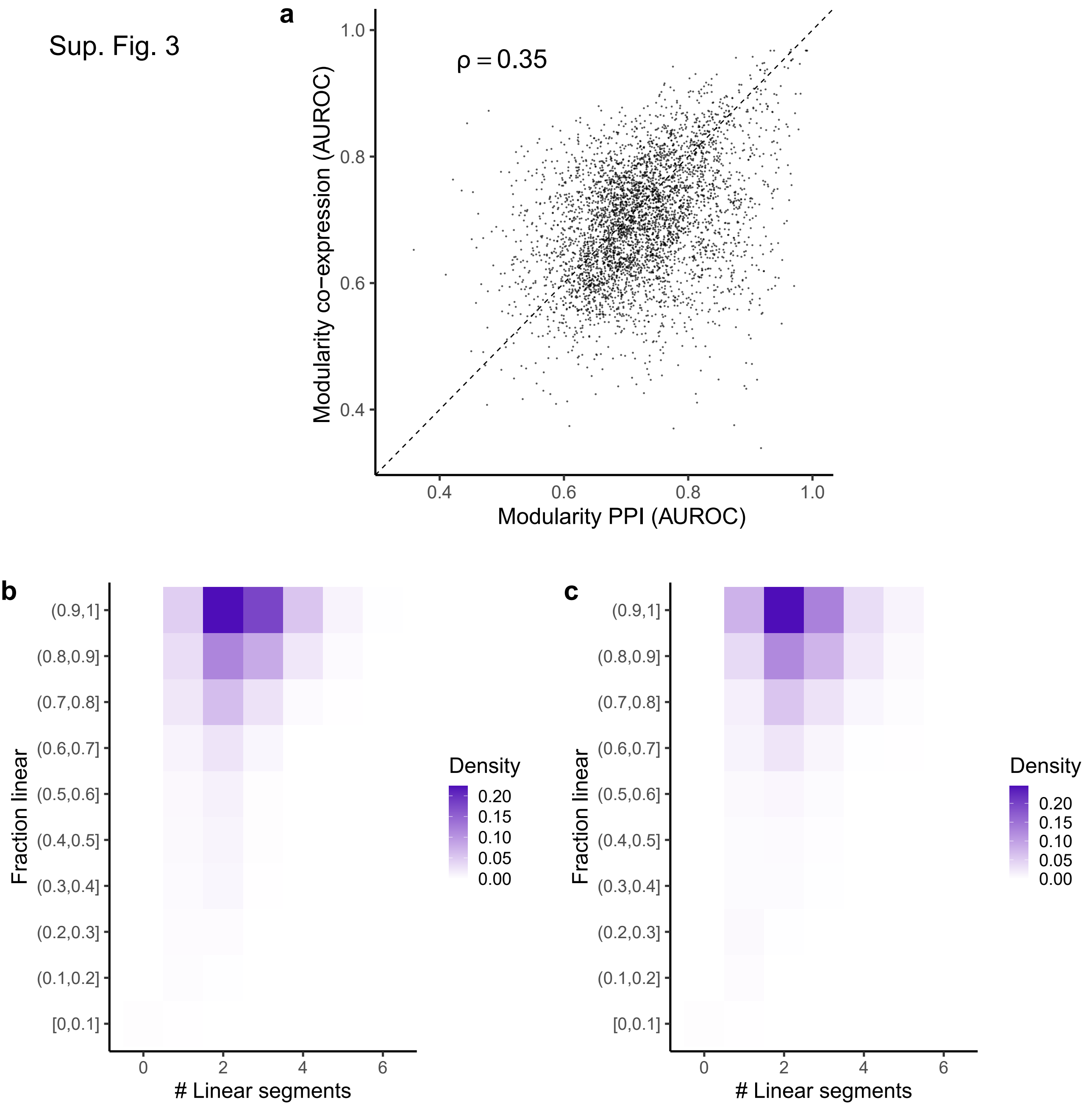

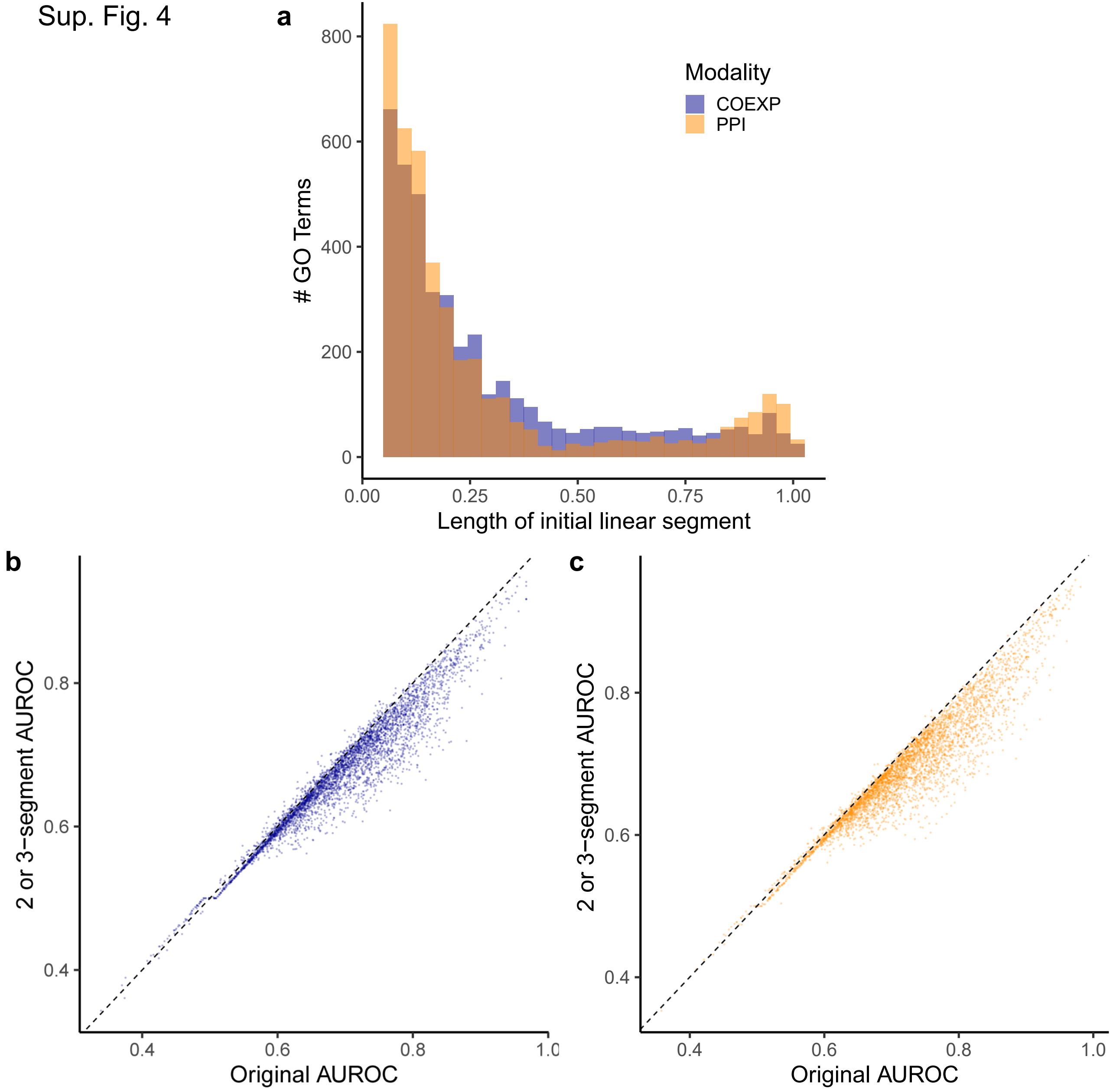

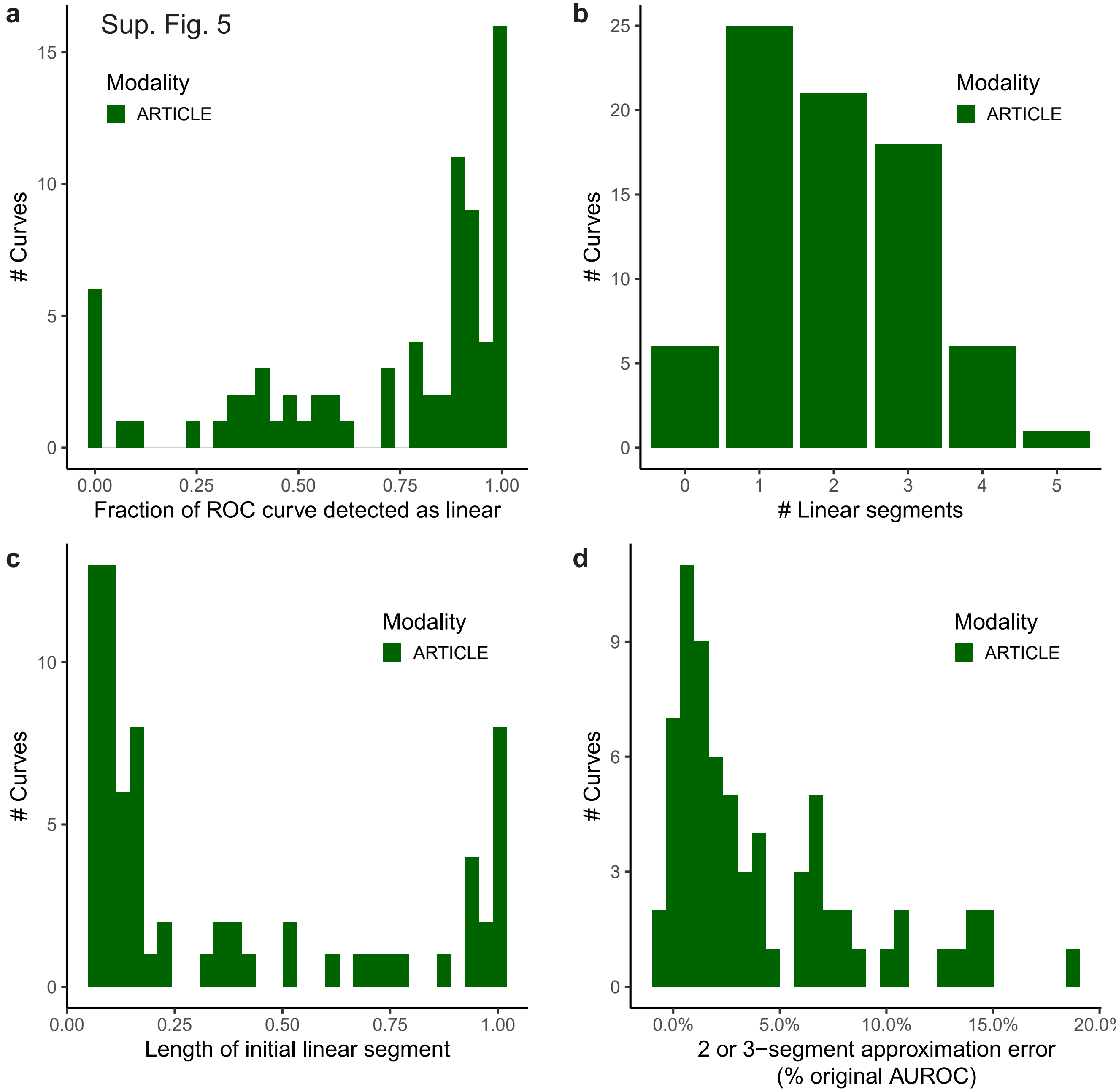
