## Supplementary Note for "Defining the extent of gene function using ROC curvature"

In our manuscript "Defining the extent of gene function using ROC curvature", we extracted 77 ROC curves from 50 research articles. First, we downloaded all research articles containing ROC curves in genomics-related Subject Areas from the PLoS One journal for one calendar year (2013-2014), leading to a selection of 35 articles (Table 1). We completed our selection with 15 manually selected high-profile publications (Table 2).

| PUBMED ID | Figure panel | Positives | Negatives | Reference |
|---|---|---|---|---|
| **24551058** | Figure S2C | ~100 | ~10000 | (3) |
| **23741529** | Figure S1A | 6000 | Effectively genome-wide choose 2 (since random) | (5) |
| **24682035** | Figure 4B | 575 | Genome-wide | (2) |
| **24743548** | Figure 2A | 190 | 2600 | (1) |
| **24621610** | Figure S1A | 50 | Effectively genome-wide (~6000) | (4) |
| **24194887** | Figure 4 | 100 | Genome-wide 6000 | (6) |
| **23977285** | Figure 2 | 250 | ~6000 (genome-wide) | (7) |
| **24498199** | Figure 5 | 2 or more | ~20000 | (8) |
| **23922946** | Figure 10A | 172 | 33 | (9) |
| **24392133** | Figure 4 | 260 | ~21K | (10) |
| **24736605** | Figure 7 | 24+ | 10K? | (11) |
| **24675610** | Figure S6 | 2000 | 8244 | (12) |
| **24349230** | Figure 5b | 1485 | 14032 | (13) |
| **24586446** | Figure 3C | 3000 | 17K? | (14) |
| **24586611** | Figure 1 | | | (15) |
| **23894279** | Figure 3 | 93 | 7000 | (16) |
| **24260261** | Figure 1 | 3638 | Random 3638 | (17) |
| **24194902** | Figure 2 | ~2K | ~10K | (18) |
| **24236095** | Figure 4E | 163 | 163 | (19) |
| **24098743** | Figure 3 | 300 | Random 1000 | (20) |
| **24489849** | Figure 5 | | | (21) |
| **24699297** | Figure 7C | | | (22) |
| **24349035** | Figure 1 | 259 | 259 choose 2 | (23) |
| **24454733** | Figure 2 | 6000 | 6000 | (24) |
| **23874989** | Figure 3 | 270 | 562*100? | (25) |
| **24194827** | Figure 1 | | | (26) |
| **24376739** | Figure 3 | 79 | 119070 | (27) |

| | | | | |
|---|---|---|---|---|
| **24391954** | Figure 7 | 35 | 86 | (28) |
| **24349449** | Figure 5 | | | (29) |
| **24475169** | Figure 3 | 93 | 93 | (30) |
| **24019945** | Figure 4 | 109 | 1700 | (31) |
| **23950912** | Figure 6A | | | (32) |
| **24069417** | Figure 6 | 4753 | 17793 | (33) |
| **23675414** | Figure 10 | | | (34) |
| **23690949** | Figure S1 | | | (35) |

**Table 1. Publications containing genomics-related ROC curves from the PLoS One journal.**

| PUBMED ID | Figure panel | Positives | Negatives | Reference |
|---|---|---|---|---|
| **18371930** | Figure 3 | 783 | | (36) |
| **24156763** | Figure S9 | 200 | ~200 | (37) |
| **16685651** | Figure 4C | 409 | | (38) |
| **22681890** | Figure S1B | | | (39) |
| **23545499** | Figure 5B | ~100 | | (40) |
| **18724933** | Figure S2C | | | (41) |
| **23932120** | Figure 2A | | | (42) |
| **20813266** | Figure S3A | 125 | | (43) |
| **16680138** | Figure 3B | 627 | | (44) |
| **24813450** | Figure 3C | 200 | | (45) |
| **20118918** | Figure 2F | ~30 | | (46) |
| **24114784** | Figure 1B | 150 | | (47) |
| **22344438** | Figure 3C | 253 | | (48) |
| **19690572** | Figure 1A | 834 | | (49) |
| **15998909** | Figure 3A | | | (50) |

**Table 2. Selection of high-profile publications containing ROC curves.**
